# Invasive terrestrial invertebrate detection in water and soil using a targeted eDNA approach

**DOI:** 10.1101/2022.11.29.518289

**Authors:** Cecilia Villacorta-Rath, Lori Lach, Natalia Andrade-Rodriguez, Damien Burrows, Dianne Gleeson, Alejandro Trujillo-González

## Abstract

1. Terrestrial invasive invertebrates can rapidly colonize new areas, causing detrimental effects on biodiversity, economy, and lifestyle. Targeted environmental DNA (eDNA) methods could constitute an early detection tool given their sensitivity to small numbers of individuals.
2. We hypothesized that terrestrial runoff would transport eDNA from the land into adjacent water bodies and used the invasive yellow crazy ant (*Anoplolepis gracilipes*) as a model species to test this hypothesis. We collected water samples from four waterbodies adjacent to infestations following rainfall events for eDNA analysis. We also collected soil samples from areas of known infestations and tested five eDNA extraction methods to determine their efficiency to extract eDNA from soil.
3. Water samples resulted in positive yellow crazy ant eDNA amplification (20–100% field replicates across all sites), even at one site located 300 m away from where ants had been detected visually. Soil samples resulted in a high percentage of false negatives when sampled from ant transit areas than from nest entrances.
4. Unpurified DNA extracts from soil also resulted in false negative detections, and only after applying a purification step of DNA extracts, we detected yellow crazy ant eDNA in 40–100% of field replicates across all methods and sites.
5. This is the first study to empirically show that eDNA from a terrestrial invertebrate can be successfully isolated and amplified from adjacent or downstream waterbodies. Our results indicate that eDNA has the potential to be a useful method for detecting terrestrial invertebrates from soil and water.

## Introduction

Over the past 15 years, environmental DNA (eDNA) analysis has gained momentum for biomonitoring of both marine and freshwater aquatic systems. Targeted-eDNA methods (i.e., qPCR-based) are now considered a sensitive, specific, and robust tool for detection of aquatic or semi-aquatic invasive species (Ficetola et al., 2008; Piaggio et al., 2014; Smart et al., 2015; Villacorta-Rath et al. 2020). This has resulted in eDNA methods being increasingly adopted into monitoring programs by natural resource management agencies, consultancy companies and citizen science groups (Darling & Mahon, 2011; Larson et al., 2020). However, advances in the use of eDNA to detect aquatic species have not been matched by developments for terrestrial species.

A key challenge in using targeted eDNA as a method to detect terrestrial species is determining an effective sampling strategy. This needs to consider where eDNA is likely to be deposited by the target organism, the effect of the substrate on eDNA detectability, and ease of sampling. To date, a few studies have used soil samples for targeted eDNA detection (Kucherenko et al., 2018; Katz et al., 2021; Yasashimoto et al., 2021) and most of them have reported false negatives arising from sampling soil areas that the species did not occupy (Kucherenko et al., 2018; Yasashimoto et al., 2021). Similarly, DNA binds to soil to a varying level depending on its physicochemical composition, and inappropriate eDNA extraction methods can lead to qPCR inhibition, affecting eDNA detectability and data interpretation (Andersen et al., 2012; Katz et al., 2021; Yasashimoto et al., 2021).

Invasive ants are among the most harmful invasive species globally (Kenis et al., 2009); they threaten the environment, human health and livelihoods, and social amenity (Holway et al., 2002; Lach & Hooper-Bùi, 2010; Gruber et al., 2022). Millions of dollars are spent on prevention, treatment and control of ant invasions globally to avoid these impacts (Zenni et al., 2021). Current methods for invasive ant detection (i.e., baited traps or cards, pitfall traps, and detection dogs) rely on luring, trapping, smelling, or sighting active individuals or colonies, which can be labour-intensive, costly, and reliant on species behaviour and weather conditions. These methods have low detection sensitivity to low numbers of individuals, which can lead to false negatives (Stringer et al., 2011, L. Lach *pers obs*.). Environmental DNA analysis could improve detectability of invasive ants, since it does not require sighting the target species (Jerde et al., 2011). However, studies investigating terrestrial insect eDNA capture for species detection are seldomly available and are designed in agricultural contexts, wherein their sampling strategy consisted of spraying water over crops to aggregate the available eDNA deposited there (Valentin et al., 2018; Valentin et al., 2020; Allen et al. 2021).

In this study we test different methods to capture and detect terrestrial invertebrate eDNA. We use the yellow crazy ant, *Anoplolepis gracilipes* (Smith, 1857), one of the most environmentally and socioeconomically damaging invasive insect species in the world (Clarke et al., 2021; Gruber et al., 2022), as model species. We collected water and soil samples in field conditions to investigate whether yellow crazy ant eDNA can be detected in water from creeks or rivers adjacent to existing infestations; and to compare the efficiency of laboratory extraction methods for eDNA extraction from soil samples.

## Materials and methods

### Study system

The yellow crazy ant is a widespread invader in tropical regions, particularly the Indo-Pacific (Janicki et al., 2016). Colonies consist of multiple interacting nests, usually termed ‘supercolonies’ (Abbott, 2005; Hoffmann, 2014). In forested areas, nests are typically at the base of trees, or under rocks, leaf litter, or logs, but the ants will nest virtually anywhere with the right temperature, humidity, and protection from sunlight (e.g., discarded car engines and soft drink cans) and can readily relocate nests when disturbed (Hoffmann, 2015; Lach, *pers. obs*.). Workers may also shelter temporarily in nest-like sites without brood or queens, for unknown periods of time (Lach, *pers. obs*.). Their variable nesting and foraging behaviour make choosing appropriate sites from which to sample soil for eDNA more challenging than it may be for other species such as Argentine ants (Yasashimoto et al., 2021). The first recorded infestation in mainland Australia was in the Northern Territory in the early 1990s (Majer, 1984). Yellow crazy ant incursions have been reported in Queensland, Australia, since 2001, including in the Cairns and Townsville regions (Lach & Hoskin, 2015).

### Environmental DNA sample collection

Water samples for eDNA analysis were collected and preserved from waterbodies adjacent to yellow crazy ant infestations in Townsville, Queensland, Australia with “high activity” (M. Green, Townsville City Council, *pers. obs*.) in February and March 2021 (Ross River, unnamed creek adjacent to Chauncy Crescent, Stuart Creek) (Table 1), during or immediately after rainfall events. We also sampled from an unnamed creek (Palmetum site) in which yellow crazy ants had been detected 300 m apart, but not adjacent (M. Green, Townsville City Council, *pers. obs*.). At each site, five replicate 30 mL surface water samples were collected using a clean 50 mL Falcon tube and decanted into another 50 mL Falcon tube containing 10 mL of Longmire’s preservative solution (Longmire & Baker, 1997) as per (Villacorta-Rath et al., 2021). Replicate water samples were taken at regular intervals, approximately 10 m apart. At every site, a field blank was also carried out to ensure that the process of sample collection did not introduce contamination. The field blank consisted of decanting 30 mL of laboratory-grade water into a Falcon tube containing 10 mL of preservative solution.

**Table 1.** Field sites where water and soil samples were collected for eDNA analyses in the Townsville area, north Queensland. All samples were collected by the authors, except for water samples from Stuart Creek, which were collected by the Townsville City Council.

| Site | Latitude (°S) | Longitude (°E) | # specimens/eDNA replicates | Collection date | Collection time | Sample volume per replicate |
| --- | --- | --- | --- | --- | --- | --- |
| **Water eDNA samples** |  |  |  |  |  |  |
| Ross River | 19.3127 | 146.7563 | 5 | 26/02/2021 | morning | 30 mL |
| Creek at Chauncy Crescent | 19.3126 | 146.7575 | 5 | 26/02/2021 | morning | 30 mL |
| Creek at Palmetum | 19.3114 | 146.7631 | 5 | 26/02/2021 | morning | 30 mL |
| Stuart Creek | 19.3405 | 146.8533 | 5 | 8/03/2021 | morning | 30 mL |
| **Soil eDNA samples - Ant transit sampling event** |  |  |  |  |  |  |
| Gieseman Road | 19.2680 | 146.5784 | 7 | 10/12/2020 | morning | 1 g |
| Copper Refinery | 19.3411 | 146.853 | 7 | 10/12/2020 | morning | 1 g |
| **Soil eDNA samples - Nest entrance sampling event** |  |  |  |  |  |  |
| Gieseman Road | 19.2680 | 146.5784 | 10 | 27/04/2021 | morning | 1 g |
| Douglas | 19.3126 | 146.7575 | 10 | 26/04/2021 | dusk | 1 g |

Soil samples were collected during two other sampling events in Townsville at known infestation sites (Table 1). Soil samples were not collected at the same time as water samples because ants tend to retreat into their nests during rainfall events, when water samples were collected. Soil sampling took place in the morning and dusk and during the wet season, when yellow crazy ant activity is likely to be high (Hoffmann, 2015). During the first sampling event, samples were collected from areas where ants could be observed transiting (hereafter referred to as ant transit samples). Seven replicate samples were collected from two sites along 50 – 70 m transects, starting where ant activity was observed. One of the sites (Gieseman Road site; Table 1) was adjacent (< 10 m away) to a water body (Stuart Creek, Table 1). Sampling consisted of collecting 2 mL of soil using a 50 mL Falcon tube and transferring 1 mL into a clean collection tube containing 700 μL cetyl trimethylammonium bromide (CTAB) buffer and 1 mL into a collection tube containing 2 mL of Biomeme lysis buffer from the M1^™^ Bulk Sample Prep Kit for DNA – High Concentration (Biomeme, Inc.). Two field controls were collected to assess potential field contamination: one was a tube containing 700 μL CTAB buffer and the other was a Biomeme homogenisation tube containing 2 mL of Biomeme lysis buffer.

In the second sampling event we collected soil from suitable microhabitats (bases of trees and under fallen logs) that had visible evidence of high yellow crazy ant traffic and high incidence of dead ants, which were considered putative nest entrances (hereafter referred to as ant nest entrance samples). In this case, ten 50 mL replicate soil samples were collected from two sites. One of the sites (Gieseman Road, Table 1) was resampled, however, the other previously sampled site (Copper Refinery) had recently been treated with insecticide, so samples were collected from a different site (Douglas: Table 1). Similar to the Gieseman Road site, the Douglas site was adjacent (< 10 m away) to a water body (Ross River, Table 1). Sampling consisted of collecting 50 mL of soil in sterile plastic containers, carefully avoiding visible dead ants. Containers were stored in ice and immediately transported to the laboratory at James Cook University. Each sample was then shaken to homogenise and approximately 1 g of soil was partitioned into four sub-samples for extraction method comparisons and stored in DNA LoBind 2 mL tubes at −80°C until eDNA was extracted. Field controls consisted of five tubes containing the lysis buffers of each eDNA extraction method (described below).

### Environmental DNA extraction from water

We extracted eDNA from water samples at the dedicated TropWATER eDNA laboratory, James Cook University, Townsville. We followed the preserve, precipitate, lyse, precipitate, purify (PPLPP) method (Edmunds & Burrows, 2020). Briefly, the PPLPP workflow uses a glycogen-aided isopropanol-based precipitation, followed by a guanidinium hydrochloride and TritonX based lysis and a subsequent glycogen-aided polyethylene glycol (PEG) based precipitation. DNA extracts were then purified to remove environmental inhibitors using the DNeasy PowerClean Pro Cleanup Kit (Qiagen^®^), as per the usual procedure of eDNA water sample analysis (Villacorta-Rath et al., 2020; Villacorta-Rath et al., 2021).

### Environmental DNA extraction from soil

Ant transit soil samples were extracted using two methods: the field-based extraction and the laboratory-based method. The field-based extraction method involved using the Biomeme M1 sample prep kit, and the laboratory-based method involved using chloroform-based extraction (CTAB; Adamkewicz & Harasewych, 1996). For the field-based extraction, 1 mL of soil sample was added to a Biomeme homogenising tube with 2 mL of Biomeme lysis buffer and shaken vigorously for one minute. After this time, the fluid was drawn from the tube using the Biomeme extraction column and syringe and pumped out 20 times. Extraction methods were then followed as per the manufacturer’s instructions and eluted in 100 μL of Biomeme Elution buffer. For the laboratory-based extraction, 1 mL of soil sample was directly transferred into a tube containing 700 μL CTAB buffer and 10 μL proteinase K was added to samples upon arrival to the laboratory following Adamkewicz and Harasewych (1996). Samples were vortexed, crushed with plastic pestles and left to lyse at 65 °C in a hibernation oven on a rocking platform for 20 hours. After lysis was performed, 700 μL chloroform-isoamyl was added and samples were centrifuged at 16,000 g for 10 min. The supernatant was then transferred into a new tube containing 600 μL chloroform-isoamyl and centrifuged again at 16,000 g for 10 min. The resulting supernatant was transferred into a new tube containing 600 μL of cold isopropanol, inverted to mix and stored at −20 °C overnight. After freezing, samples were centrifuged at 16,000 g and at 4 °C for a total of one hour and the supernatant was pipetted off taking care to not lose the formed pellet. The pellet was then washed with 1,000 μL of 70% ethanol and centrifuged at 16,000 g for 10 min. Finally, the ethanol was pipetted off and pellets were allowed to air dry for 15 min, before resuspending them in 100 μL TE buffer and stored at 4 °C. DNA extracts from both extraction methods were tested for presence of contaminants using a NanoDrop^™^ spectrophotometer.

Ant nest entrance soil samples were extracted using five methods: (1) the Biomeme M1 sample prep kit; (2) the CTAB method; (3) the PPLPP method (Edmunds & Burrows, 2020); (4) the Qiagen^®^ DNeasy PowerSoil kit; and (5) the modular-universal DNA extraction (Mu-DNA) method (Sellers et al. 2018).

For the Biomeme M1 sample prep kit, we modified the previously described procedure in the first and last steps: 1 mL of soil samples were initially transferred into 3 mL of Biomeme lysis buffer, and DNA extracts were eluted in 400 μL Biomeme Elution buffer. Similarly, the first step of the CTAB method was modified, wherein each 1 mL-field replicate was split into two 2 mL DNA LoBind^®^ tubes (approximately 0.5 mL of soil sample/tube) containing 1000 μL CTAB buffer.

For the PPLPP method, each replicate of 1 mL soil sample was transferred into a 50 mL DNA LoBind^®^ Falcon tube containing 10 mL Longmire’s buffer (Longmire & Baker, 1997) and 10 mL of MilliQ water. Environmental DNA was extracted following Edmunds and Burrows (2020) with eDNA eluted in 100 μL.

For the Qiagen^®^ DNeasy PowerSoil kit, each field replicate consisting of 1 mL soil was partitioned into four tubes with 250 mL of soil and mixed with 60 μL Solution C1. The bead beating step was not performed given the target was not bacterial DNA from the soil samples. We then followed the manufacturer’s protocol handling each field replicate in four separate tubes, sequentially passed through a single spin column and eluted in 100 μL elution buffer.

Finally, we followed the soil sample workflow of the Mu-DNA protocol without the bead beating step. Each 1 mL soil replicate was split into two 2 mL DNA LoBind^®^ tubes and mixed with 550 μL lysis solution, 200 μL soil lysis additive and 20 μL proteinase K. Samples were then vortexed and incubated for 3 h at 55°C. Subsequently, samples were centrifuged at 4,000 g for 1 min, the supernatant was transferred into a new tube, centrifuged at 10,000 g for 1 min and the supernatant was again transferred into a new tube containing 300 μL flocculant solution. Extraction was then carried out as published (Sellers et al. 2018). A negative extraction control was added to each batch of eDNA extractions to ensure that no contamination was introduced during laboratory procedures.

### Quantitative real-time polymerase chain reaction (qPCR)

We screened samples for yellow crazy ant eDNA presence using two probe-based, species-specific eDNA assays developed and optimized by EcoDNA, targeting two separate sections of the yellow crazy ant Cytochrome Oxidase 1 (COI) gene region: Agra1 assay (112 base pair [bp] long) and Agra2 assay (131 bp long) (Supporting Information 1).

qPCR plates were set-up using the EzMate^™^ 401 Automated Pipetting System (Arise Biotech) and run in a QuantStudio^™^ 5 Real-Time PCR System (Thermo Fisher Scientific Australia Pty Ltd). We tested four technical replicates of each sample and each site, including field and extraction blanks, three no-template controls, and genomic DNA positive controls. Each qPCR reaction and cycling conditions were as explained in the assay development section of this study. Inhibition was tested using a TaqMan^™^ Exogenous Internal Positive Control (IPC) qPCR assay (ThermoFisher Scientific). A total of 3 μL IPC was applied to duplicate samples and three reactions containing only IPC were included as controls. A departure of three or more cycles (ΔCt) would indicate sample inhibition (Hartman et al., 2005). Samples were subsequently purified using the DNeasy PowerClean Pro Cleanup Kit (Qiagen^®^) and another qPCR was carried out. Ct values between unpurified and purified samples were compared to assess the level of sample inhibition. A subset of amplicons with positive detections were Sanger sequenced for confirmation of results at the Australian Genome Research Facility (AGRF).

### Data analysis

Differences in yellow crazy ant eDNA capture sensitivity (number of DNA copies per assay) across different methods were assessed with a generalized linear mixed model using a template model builder (TMB) computed in the R package glmmTMB version 1.7.19 (Brooks et al., 2021). The response variable was DNA copy number, and the explanatory variables were eDNA extraction method (fixed effect) and field replicate/technical replicate (nested random effects, with technical replicates nested within field replicates). Two models were run: the first one testing replicates as an additive fixed factor and the second one testing replicates as an interaction. The best performing model was chosen based on the corrected Akaike information criterion (AICc). We tested for overdispersion with the DHARMa R package version 0.4.4 (Hartig, 2021). Post-hoc paired comparisons of means were performed using Tukey’s HSD. Statistical analyses were completed in R (R Development Core Team, 2021).

## Results

### Yellow crazy ant eDNA detection in water and soil

Water samples collected adjacent or in the vicinity of yellow crazy ant infestations showed positive eDNA amplification with both assays. The highest percentage of eDNA detections were observed at Stuart Creek (100% of field and technical replicates using the Agra2 assay), followed by Ross River (80% and 60% of field and technical replicates using the Agra2 assay, respectively), Palmetum creek (100% and 50% of field and technical replicates using the Agra2 assay, respectively), and Chauncy Crescent creek (20% and 10% of field and technical replicates using the Agra2 assay, respectively) (Fig. 1). The Agra2 eDNA assay amplified DNA extracts from water samples at a higher percentage than the Agra1 assay, which failed to detect yellow crazy ant eDNA at Chauncy Crescent creek (Fig. 1).

**Figure 1.**
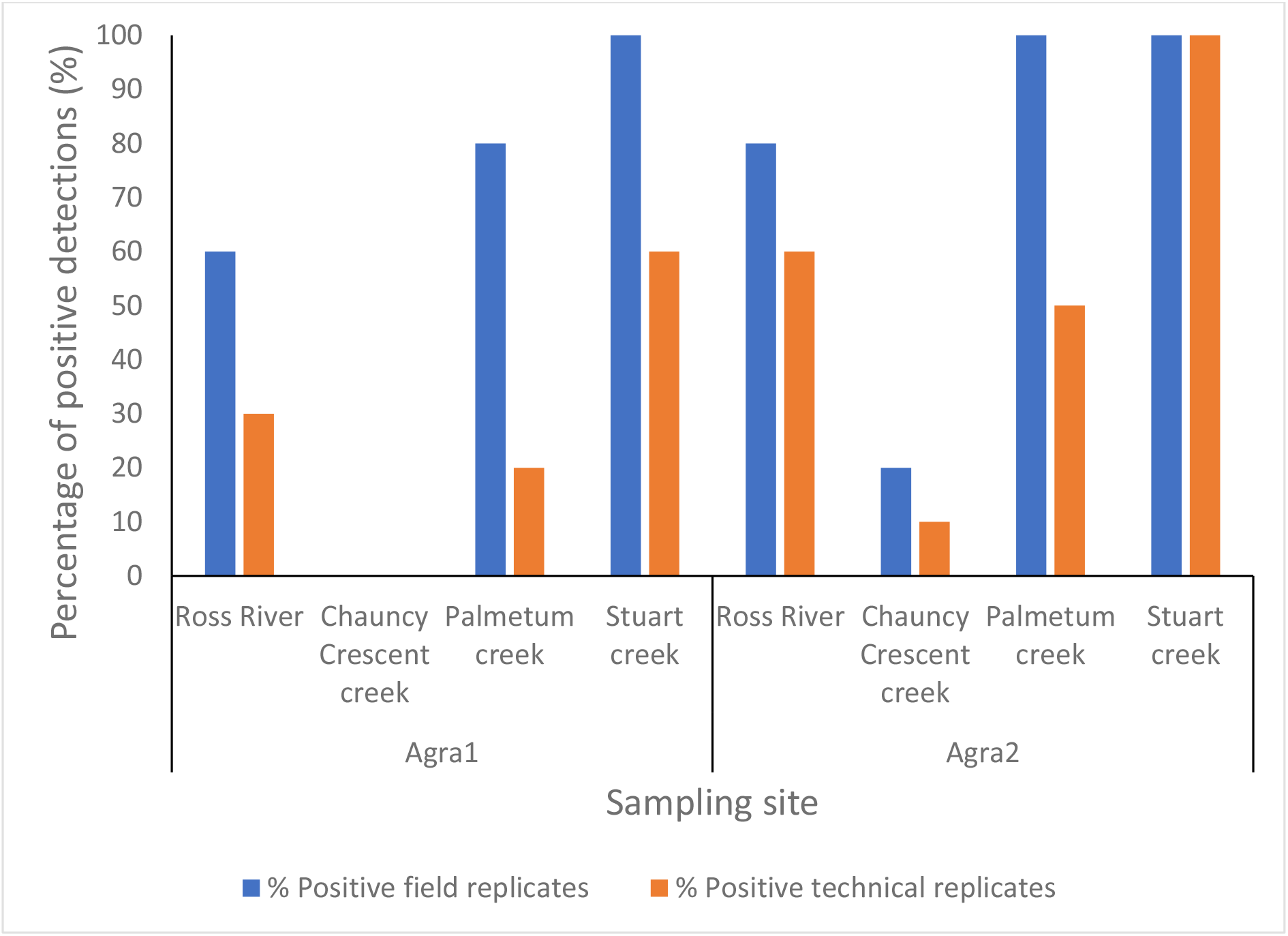
Percentage of positive yellow crazy ant eDNA detections from purified water samples using the Agra1 and Agra2 eDNA assays, targeting two different fragments of the COI gene.

In soil samples, the Agra1 assay was more efficient at amplifying DNA extracts. Samples collected from ant transit sites showed high concentration of contaminants (Table S1), and complete inhibition was observed in eDNA samples tested using both eDNA assays (Table S2). After purification of DNA extracts, both assays successfully amplified 43-64% of the laboratory-extracted samples (Tables 2, S3). However, *in-situ* extracted samples had a low percentage of positive detections (14-18%) (Tables 2, S3).

**Table 2.**
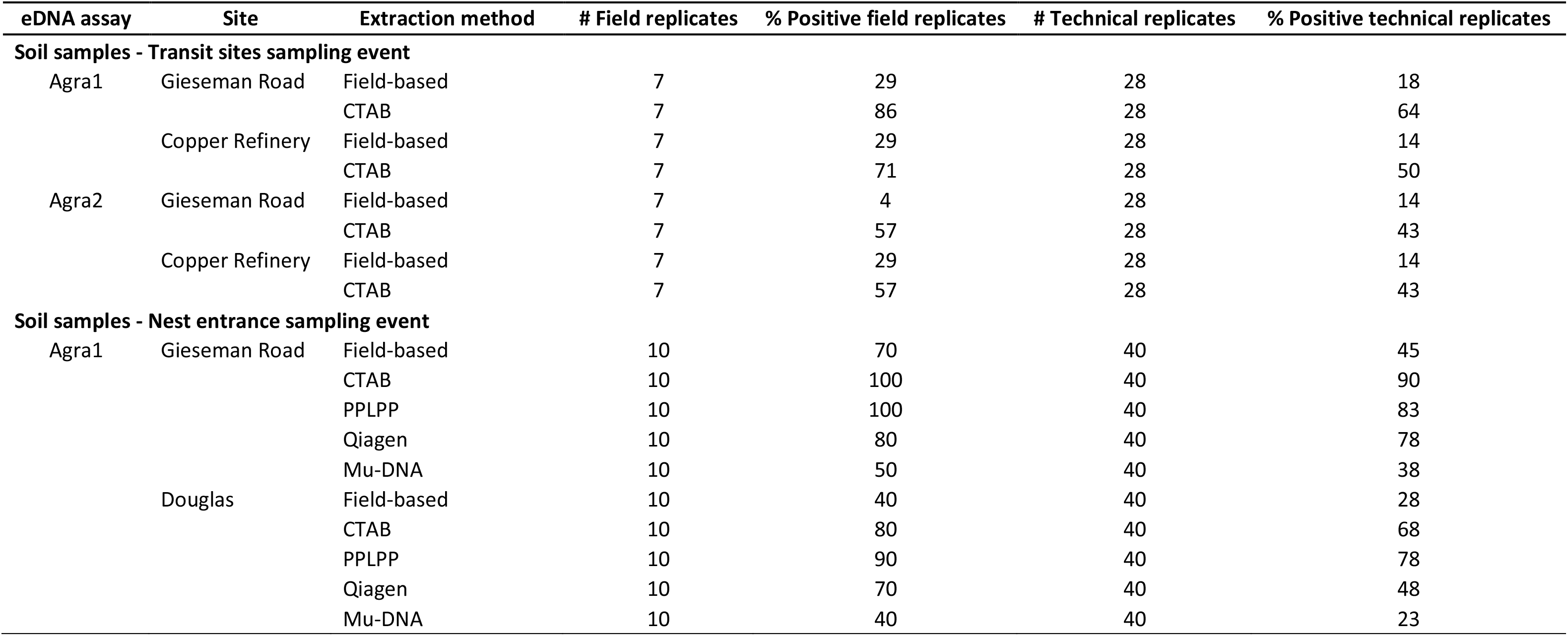
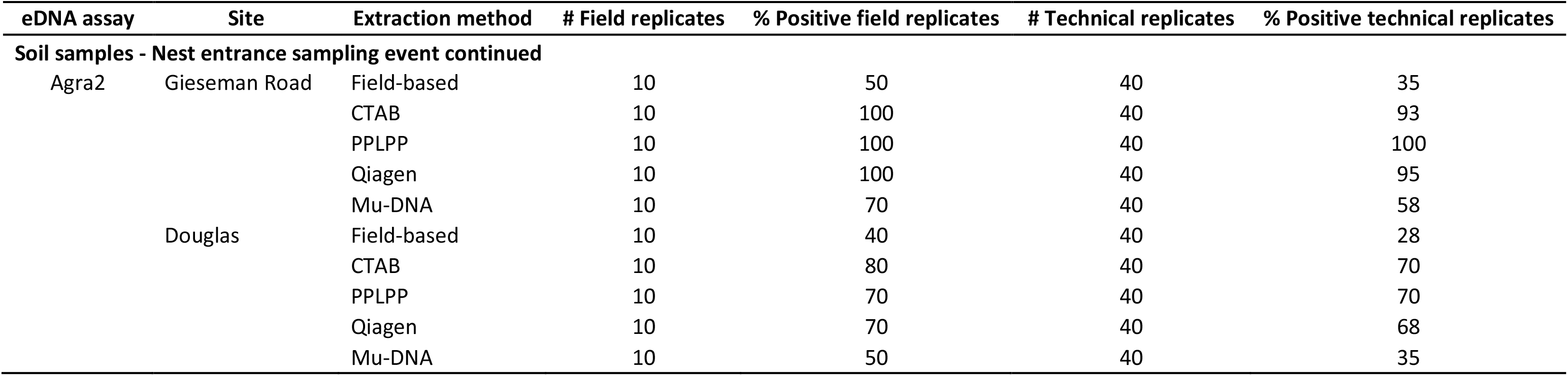
Yellow crazy ant eDNA detection in purified soil samples using Agra1 and Agra2 eDNA assays, targeting two different fragments of the COI gene. Extraction methods used were field-based (Biomeme^®^ M1 sample prep kit), CTAB (cetyltrimethylammonium bromide, Adamkewicz and Harasewych, 1996), PPLPP (preserve, precipitate, lyse, precipitate, purify method, Edmunds & Burrows, 2020), Qiagen (Qiagen^®^ DNeasy PowerSoil kit), and Mu-DNA (modular-universal DNA extraction method, Sellers et al. 2018).

There was a greater percentage of positive technical replications from nest entrance samples compared to transit samples for both assays and for field based and CTAB purification (Tables 2, S4). In general, Agra1 assay was more efficient at amplifying yellow crazy ant eDNA, reflected in higher eDNA yields across all extraction methods than Agra 2 assay (Table S4). Also, the CTAB, PPLPP and Qiagen extraction methods had the highest percentage of positive detections when using both assays (Table S4), whereas eDNA yield (number DNA copies/assay) was variable depending on the sampling site and eDNA assay used (Fig. 2, Table S4). Field and extraction controls from both the ant transit sites and nest entrance sampling events did not show positive amplification.

**Figure 2.**
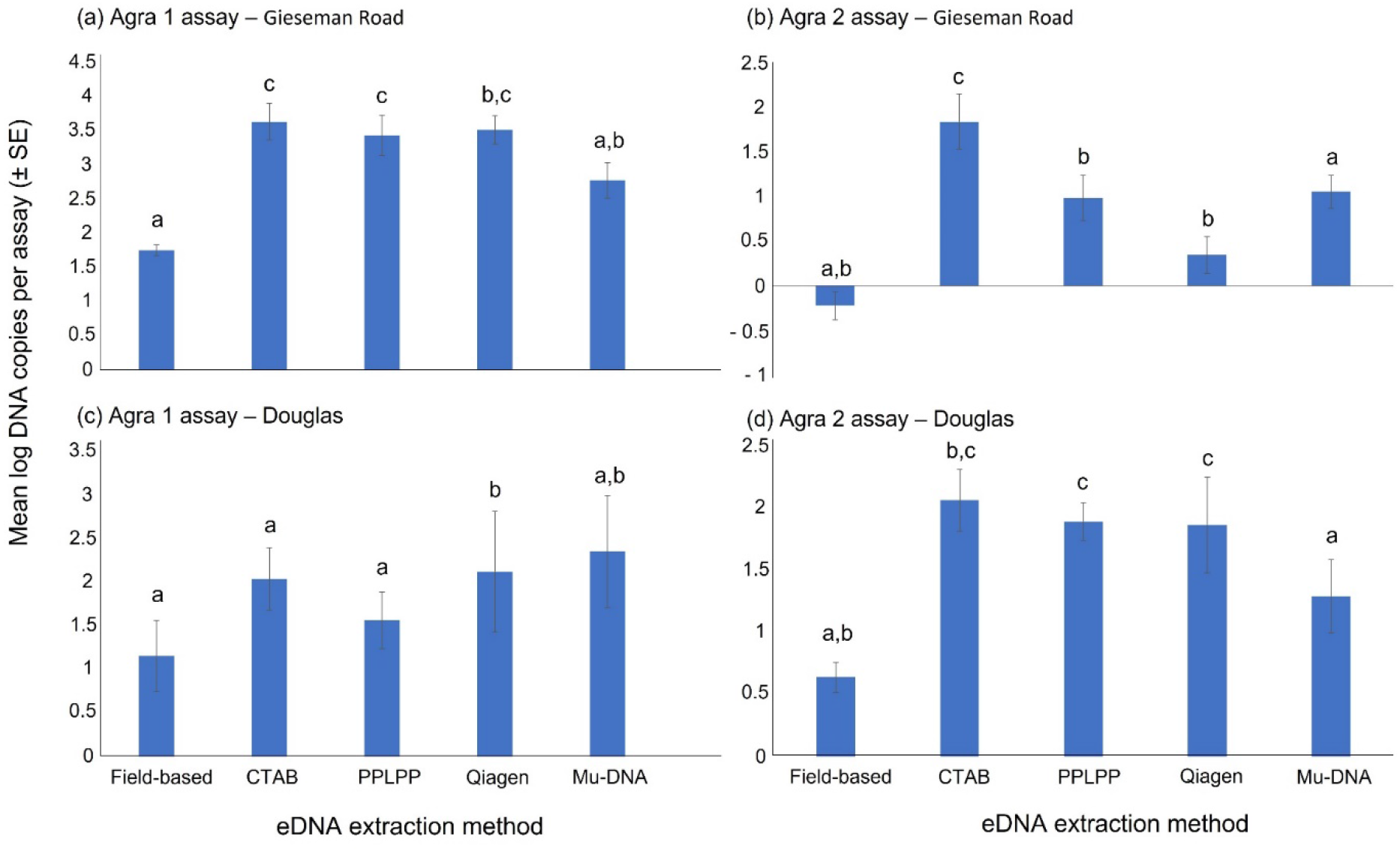
Mean DNA concentration (mean DNA copy number per assay ± Standard Error) yielded by each of the five eDNA extraction methods from purified soil samples collected from likely yellow crazy ant nest entrances (ant nest entrance sampling event) and run using the (a) Agra1 assay on Gieseman Road samples, (b) Agra2 assay on Gieseman Road samples, (c) Agra1 assay on Douglas samples, and (d) Agra2 assay on Douglas samples. DNA yield was log10 transformed. Note the differences in y-axis scales across panels. Methods with different letters above the bars within each panel differ significantly in post-hoc tests (at *P* < 0.05) using Tukey HSD. Extraction methods used were field-based (Biomeme^®^ M1 sample prep kit), CTAB (cetyltrimethylammonium bromide, Adamkewicz and Harasewych, 1996), PPLPP (preserve, precipitate, lyse, precipitate, purify method, Edmunds & Burrows, 2020), Qiagen (Qiagen^®^ DNeasy PowerSoil kit), and Mu-DNA (modular-universal DNA extraction method, Sellers et al. 2018).

At Ross River, where we collected both water and soil samples, yellow crazy ant eDNA detections of both substrata were similar: 40-90% of positive soil field replicates (Agra1 assay) compared to 80% of positive water field replicates (Agra2 assay), and 28-78% of positive soil technical replicates (Agra1 assay) compared to 60% of positive water field replicates (Agra2 assay).

### Comparison of eDNA yield across extraction methods

There were significant differences between mean DNA copy number across all eDNA extraction methods at both sites and using both assays (Fig. 2). For Gieseman Road samples tested using Agra1 assay, the field-based method yielded significantly fewer DNA copies than the CTAB (F = −3.006, *P* = 0.0008), PPLPP (F = −3.888, *P* < 0.001) and Qiagen (F = −2.345, *P* = 0.0136) methods (Fig. 2). Also, the Mu-DNA method yielded significant fewer DNA copies than the CTAB (F = 2.198, *P* = 0.0345) and PPLPP (F = −3.080, *P* = 0.0008) methods (Fig. 2). When using the Agra2 assay we found that the CTAB method yielded significantly higher number of DNA copies than the field-based (F = −2.590, *P* = 0.0018), Mu-DNA (F = 4.295, *P* < 0.0001), PPLPP (F = 1.711, *P* = 0.0003) and Qiagen (F = 1.886, *P* = 0.0001) methods (Fig. 2). Also, the Mu-DNA method had significantly higher number of DNA copies than the PPLPP (F = −2.583, *P* = 0.0001) and Qiagen (F = −2.409, *P* = 0.0004) methods (Fig. 2).

For Douglas samples, the Qiagen method exhibited significantly higher number of DNA copies than the field-based (F = −7.477, *P* = 0.0002), CTAB (F = −4.399, *P* = 0.0115) and PPLPP (F = −6.051, *P* < 0.0001) methods of samples tested using the Agra1 assay (Fig. 2). When using the Agra2 assay, the field-based method yielded significantly fewer DNA copies than the PPLPP (F = −2.8819, *P* = 0.0036) and Qiagen (F = −3.5810, *P* = 0.0118) methods (Fig. 2); and the Mu-DNA method had significantly fewer DNA copies than the CTAB (F = 2.4420, *P* = 0.0077), PPLPP (F = −2.9663, *P* = 0.0039) and Qiagen (F = −3.6653, *P* = 0.0002) methods (Fig. 2).

## Discussion

Detection methods that are sensitive to small number of individuals, such as eDNA analysis, have the capacity to complement and improve the detection of invasive species. We used yellow crazy ants as a model species to test eDNA detection of a terrestrial invasive species in water and soil, as well as to explore the effect of soil eDNA extraction methods on eDNA detectability. Both substrata yielded positive yellow crazy ant eDNA detection. Importantly, we detected yellow crazy ant eDNA in water samples from creeks and rivers directly adjacent and in the vicinity of known infestations. To the best of our knowledge, our findings are the first demonstration of the feasibility of detecting terrestrial invertebrate eDNA in natural waterways. Additionally, we found that eDNA detectability in soil is dependent on the extraction method and the area from which the samples are collected (i.e., ant transit areas vs. nest entrances), and that purification of DNA extracts is necessary to avoid false negative detections.

Many aquatic eDNA studies show that population size (Yates et al., 2019; Spear et al., 2021; Yates et al., 2021), time elapsed since a target organism has occupied a system (Kucherenko et al., 2018; Schmidt et al., 2021) and target species behavior (Buxton et al., 2017; Dunn et al., 2017) influence eDNA detectability in water. In the present study, qualitative data from the infestations adjacent to our water sampling sites suggested that ant activity (behaviour) was high everywhere except next to Palmetum creek, where 80% of the field replicates exhibited yellow crazy ant eDNA. On the other hand, at Chauncy Crescent creek, adjacent to an area of high ant activity, only 20% of the field replicates resulted in positive eDNA detections. Therefore, future research to disentangle the factors related to ant density and activity that could affect eDNA detectability in water will be useful. It is also expected that the amount of rainfall prior to water sampling and the distance between the infestations and receiving waterbodies will play an important role in transporting terrestrial eDNA into the aquatic system. Once yellow crazy ant eDNA is in the aquatic system, we hypothesize that the amount of rainfall will influence eDNA detectability by increasing water flow and dilution. As with aquatic eDNA studies, we would expect that the time elapsed since eDNA transport into aquatic systems would determine eDNA detectability due to factors affecting aquatic eDNA detection (i.e., eDNA production, decay, transport, retention, and resuspension; Barnes et al., 2016). Future studies should focus on investigating four main factors that could influence yellow crazy ant eDNA detectability in waterbodies, namely: (1) total area of the infestation; (2) ant activity; (3) time since the establishment at a site; and (4) amount of rainfall prior to water sampling.

In soil samples, eDNA detectability from areas of ant transit was lower than that of ant nesting areas. A recent study on Argentine ants eDNA also found the highest eDNA concentration in soil from nest entrances, as opposed to surface soil samples from an infestation area and found no relationship between eDNA concentration and distance from nests or trails (Yasashimoto et al., 2021). The authors argued that Argentine ants may move nests frequently, and therefore strong relationships between eDNA concentration and distance from a nest were not expected (Yasashimoto et al., 2021). Yellow crazy ants also move nests and transfer brood to different locations frequently (Lach *pers. obs*.), and as with most ants, move dead ants to immediately outside of the nests. Therefore, it is at the nest entrances where we would expect a significant amount of eDNA to be deposited. There is also the possibility that higher detections at nest entrance sites are due to small ant parts present in soil samples, even though we avoided sampling dead ants. If eDNA methods are used to check the progress of eradication efforts, this could constitute a source of false positive detections. Therefore, it would be important to investigate how to avoid the potential of false positive detections arising from dead yellow crazy ants. If the aim is to detect presence of the species in a new area, we propose the highest likelihood of collecting yellow crazy ant eDNA in soil is from samples at the base of trees or other moist areas where they are more likely to establish long-term nests. Yasashimoto et al. (2021) also concluded that the type of ant activity and their behaviour at different areas will determine eDNA detectability, indicating the importance of understanding the ecology of the species to avoid false negative detections.

Soil type may have also affected detectability. Samples collected from Gieseman Road, which has coarse sandy soils (Murtha 1975), showed a higher percentage of positive detections than those from Douglas, which has clay soils (Murtha 1975). Regardless of the eDNA assay used, soils with higher percentage of organic matter or clay and higher pH (i.e., more negatively charged) are more likely to bind to eDNA (Allemand et al., 1997; Andersen et al., 2012) and therefore inhibit the qPCR reaction. Therefore, we would expect to have more effective eDNA extraction from the coarser soil from Gieseman Road compared to the more organic-rich soil from Douglas (Murtha, 1975), which is shown by the higher percentage of positive eDNA detections found at the former.

Our results showed that column-based eDNA extraction methods (Qiagen and Mu-DNA) perform better at removing sample inhibition than the other three methods, which only showed eDNA amplification after a purification step. This means that the purification step could be avoided, cutting laboratory costs and shortening the sample processing time. In terms of eDNA yield, the Qiagen method was more or equally as effective in recovering eDNA from soil than the CTAB and PPLPP. Although the Mu-DNA method was less efficient than Qiagen, it can be scaled up to any starting volume of soil and it is almost ten times more cost-effective than the DNeasy PowerSoil kit (Qiagen^®^) (Sellers et al., 2018).

## Conclusions

In the present study we used yellow crazy ants as a model species to explore eDNA detectability in two different substrata: water and soil. We demonstrated that terrestrial eDNA can be detected in water bodies near yellow crazy ant infestations. Our findings suggest there are opportunities for detecting terrestrial invertebrate eDNA across large areas given that mechanisms such as rainfall runoff could aggregate eDNA into nearby or downstream waterbodies. However, the factors influencing terrestrial invertebrate eDNA detectability in water should be explored further. We showed that detectability of eDNA in soil is dependent on sampling location, and eDNA extraction method, and that purification of DNA extracts is important to avoid false negative detections, making soil sampling less attractive than water sampling.

## Authors’ contributions

C.V.R., A.T.G., D.G., D.B., and L.L. conceived the ideas and designed methodology; C.V.R., A.T.G. and N.A.R. collected the data; C.V.R., A.T.G., and L.L. analysed the data; C.V.R. and A.T.G. led the writing of the manuscript. All authors contributed critically to the drafts and gave final approval for publication.

## Acknowledgements

We acknowledge the Traditional Owners of the land where the sampling and laboratory analyses were conducted. The contribution of C.V.R. and N.A.R. were supported through funding from the Australian Government’s National Environmental Science Program—Northern Australia Environmental Resources Hub, Project 4.3 (to D.B.) and the Department of Agriculture, Water and Environment (to C.V.R. and L.L.). Also, the contribution of A.T.G. was funded by the Biosecurity Innovation Program, from the Australian Government Department of Agriculture, Water and the Environment (to D.G. and A.T.G.). Thanks to Angela Strain (JCU) and the Wet tropics Management Authority (WTMA) for providing yellow crazy ant specimens for assay development. We thank Melissa Green (Townsville City Council), as well as Bev Job and Janet Cross (Invasive Species Council) for assistance with sample collection and discussions on locations of yellow crazy ant infestations in Townsville.

## Data accessibility

All data supporting the findings of this study can be found under the Supporting Information and archived in the James Cook University Research Data Hub.

