## Supplemental Information 1 for "Invasive terrestrial invertebrate detection in water and soil using a targeted eDNA approach"

### Methods

*Specimen collection for eDNA assay development*

We collected five yellow crazy ant workers from each of five infestations that were at least 5 km apart in the Cairns, Queensland, Australia region (Table 1). Ant genomic DNA (gDNA) was extracted at the Molecular Ecology and Evolution Laboratory (MEEL, James Cook University) using the cetyl trimethylammonium bromide (CTAB) method (Adamkewicz & Harasewych, 1996).

**Table 1.** Yellow crazy ant specimens collected from five separate incursions in Cairns, northeast Queensland, used for eDNA assay development and optimization

| Infestation site | Latitude (°) | Longitude (°) | # specimens | Field collection date |
|---|---|---|---|---|
| Vico Street | -17.126057 | 145.757797 | 5 | 14/07/2020 |
| Wrights Creek 3 | -17.080637 | 145.723643 | 5 | 26/08/2020 |
| Henley's Hill | -16.953257 | 145.735828 | 5 | 25/03/2020 |
| Green Forest | -16.816716 | 145.576750 | 5 | 29/05/2019 |
| Gordonvale | -17.085828 | 145.794929 | 5 | 11/03/2020; 10/04/2020; 08/05/2020 |

*Assay development and optimization*

We designed two probe-based assays to amplify two separate sections of the yellow crazy ant Cytochrome Oxidase 1 (COI) gene region (Table 2) by aligning accessioned yellow crazy ant sequences in Geneious R10. Both assays were *in-silico* tested for specificity using the BLAST search function on the National Centre for Biotechnology Information (NCBI) website and checked for quality using

NetPrimer (Primer Biosoft). Each reaction consisted of 8 µL MilliQ® water, 10 µL TaqMan® Environmental Master Mix 2.0 (ThermoFisher), 1 µL of each of forward and reverse primers (10 µM), 1 µL probe (10 µM) and 5 µL DNA template for a total volume of 25 µL. Cycling conditions for both assays were: 95 °C (10 min), followed by 50 cycles of 95 °C (20 s), 60 °C (40 sec) ramping at 2.42 °C/sec, followed by a final holding stage at 4 °C in a Viia™ 7 Real-Time PCR System (Applied Biosystems, Australia).

Assays were tested for specificity with genomic DNA (gDNA) extracts of ant species provided by the Operational Science & Surveillance group in Biosecurity Operations Division of DAWE (*Anoplolepis custodiens* (Smith, 1858), *Lepisiota frauenfeldi* (Mayr, 1855), *Nylanderia fulva* (Mayr, 1862), *Solenopsis geminata* (Fabricius, 1804), *Solenopsis invicta* (Buren, 1972), *Paratrechina longicornis* (Latreille, 1802), and *Wasmannia auropunctata* (Roger, 1863). In the same way, qPCR efficiency and sensitivity were assessed by obtaining the limit of quantification (LOQ) and limit of detection (LOD) using gBlocks synthetic oligo standards designed for each assay based on the targeted fragment and *A. gracilipes* genomic DNA. Standard curves were established using dilution series of known concentrations ranging from $10^7$ copies/µL and decreasing tenfold down to 1 copy/µL, for a total of six replicates per dilution.

**Table 2.** Species-specific TaqMan® assays designed to detect yellow crazy ant Cytochrome Oxidase 1 gene region

| Assay name | Oligo | Melt temp (°C) | GC content (%) | Amplicon fragment size | Nucleotide sequence (5′–3′) |
|---|---|---|---|---|---|
| Agra1 | 566F | 60.1 | 47.8 | | CAGCAATTCTCCTCCTTCTGTCT |
| | 677R | 62.5 | 66.7 | | GGATCGCCCCCACCTGATG |
| | 589P | 63.1 | 52.2 | 112 | JUN-CTCCCAGTTTTAGCCGGAGCAAT-QSY |
| Agra2 | 793F | 60 | 34.5 | | ACTTTTGGTGCTTTAGGAATAATCTATGC |
| | 923R | 60.1 | 33.3 | | ATTGTTGCAGAAGTAAAATAAGCTCGA |
| | 843P | 62.5 | 42.3 | 131 | FAM-AGGCTTTATTGTTTGGGCTCATCACA-QSY |

**Results**

*Environmental DNA assay specificity and sensitivity*

Agra1 and Agra2 assays successfully amplified two separate 112 and 132 bp fragments, respectively, of the CO1 gene region of the targeted species, using gDNA from Cairns colonies and eDNA from Townsville colonies. No amplification was observed in DNA extracts from *A. custodiens*, *L. frauenfeldi*, *N. fulva*, *S. geminata*, *S. invicta*, *P. longicornis* or *W. auropunctata*. The LOD for the Agra1 assay was estimated to be 1 copy/µL (mean $C_T$ ± SD = 39.65 ± 0.82) with an LOQ of 1000 copies/µL (mean $C_T$ ± SD= 35.075 ± 0.32) with an $R^2$=0.95 (Table 3). Similarly, the LOD for the Agra2 assay was estimated to be 100 copies/µL (mean $C_T$ ± SD = 40.396) with an LOQ of 1000 copies/µL (mean $C_T$ ± SD=

37.223 ± 0.343) with an $R^2$=0.997 (Table 3). Amplification efficiencies ranged from 76-86% for Agra1 and 81-94% for Agra2 in plates assayed in this study. Positive controls amplified in all plates and no amplification occurred in the negative template controls.

**Table 3.** Checklist of Minimum Information for Publication of Quantitative Real-Time PCR Experiments (MIQE) for Agra1 and Agra2 eDNA assays. Essential and desirable information are included

| Item to check | | Agra1 eDNA assay |
|---|---|---|
| Experimental design | Definition of experimental and control groups | Experimental: (1) genomic DNA (gDNA) from the target species |
| | | (2) environmental DNA (eDNA) samples consisting of soil and water from known infestation areas |
| | | Control: MilliQ water |
| | Number within each group | gDNA: n = 25; eDNA: n soil = 34; eDNA: n water = 20 |
| Sample (gDNA) | Description | Water samples were collected from four creeks/rivers adjacent to yellow crazy ant incursions. Soil samples were collected from three sites of known incursions. Tissue was sourced from five separate incursions in Cairns |
| | Microdissection or macrodissection | DNA from whole yellow crazy ant specimens was extracted |
| | Processing procedure:<br>If fixed – with what, how quickly? | Tissue samples were preserved immediately in 95% ethanol |
| | Sample storage condition and duration (especially for FFPE samples) | Extracted DNA was stored at 4°C |
| Sample (eDNA) | Description | Water samples: Direct collection and preservation of 30 mL water from water body adjacent to yellow crazy ant incursions |
| | | Soil samples: Direct collection of 1mL soil from sites of known incursions kept in ice for 2 hours and then extracted in the laboratory |
| | Processing procedure | Water samples were extracted using the PPLPP method; soil samples were extracted using the M1™ Bulk Sample Prep Kit for DNA – High Concentration (Biomeme, Inc), CTAB, PPLPP, Qiagen and Mu-DNA methods |
| | Sample storage condition and duration (especially for FFPE samples) | Extracted DNA was stored at 4°C for one week (until qPCR was performed) and subsequently stored at -20°C |
| Nucleic acid extraction (gDNA) | Procedure and/or instrumentation:<br>Name of kit and details of any modifications | Cetyltrimethylammonium bromide (CTAB) method |

| | | |
|---|---|---|
| | Details of DNase or RNase | N/A |
| | Contamination assessment (DNA or RNA) | Nanodrop |
| | Nucleic acid quantification | |
| | Instrument and method | Quantus ONE kit |
| | RNA integrity method/instrument | |
| | RIN/RQI or Cq of 3' and 5' transcripts | N/A |
| | Inhibition testing (Cq dilutions, spike or other) | N/A |
| Nucleic acid extraction (eDNA) | Procedure and/or instrumentation | |
| | Name of kit and details of any modifications | eDNA extarction methods: (1) PPLPP; (2) M1™ Bulk Sample Prep Kit for DNA – High Concentration, Biomeme, Inc; (3) CTAB; (4) modular-universal DNA extraction (Mu-DNA); (5) Qiagen® DNeasy PowerSoil kit. There was a subsequent purification using the DNeasy PowerClean CleanUp Kit (Qiagen®) |
| | Details of DNase or RNase | N/A |
| | Contamination assessment (DNA or RNA) | Nanodrop |
| | Nucleic acid quantification | |
| | Instrument and method | N/A |
| | RNA integrity method/instrument | |
| | RIN/RQI or Cq of 3' and 5' transcripts | N/A |
| | Inhibition testing (Cq dilutions, spike or other) | |
| qPCR target information | Gene symbol | N/A |
| | Sequence accession number | NCBI MZ330820-MZ330832 |
| | Amplicon length | 112 bp |

| | | |
|---|---|---|
| | *In silico* specificity Screen (BLAST, etc.) | BLAST |
| | Location of each primer and probe by exon or intron (if applicable) | |
| | What splice variants are targeted? | N/A |
| | Primer and probe sequences | Forward primer: 5'–CAGCAATTCTCCTCCTTCTGTCT–3'<br>Reverse primer: 5'–GGATCGCCCCCACCTGATG–3'<br>Probe: 5'–JUN-CTCCCAGTTTTAGCCGGAGCAAT-QSY–3' |
| | Location and identity of any modifications | Cytochrome Oxidase 1 (COI) region of the mitochondrial genome |
| | Complete reaction conditions: | |
| | Reaction volume and amount of cDNA/DNA | Reaction volume: 25 µL; amount of DNA: 5 µL |
| qPCR oligonucleotides | Primer, probe, $Mg^{2+}$ and dNTP concentrations | Each 25 µL reaction contained: 10 µL Environmental master Mix 2.0; 1 µL forward primer (10 µM); 1 µL reverse primer (10 µM); 1 µL probe (10 µM); 7 µL MilliQ water; 5 µL template DNA |
| | Polymerase identity and concentration | Applied Biosystems® TaqMan® Environmental Master Mix 2.0 (Thermo Fisher Scientific) |
| qPCR protocol | Buffer/kit identity and manufacturer | Applied Biosystems® TaqMan® Environmental Master Mix 2.0 (Thermo Fisher Scientific) |
| | Additives (SYBR Green I, DMSO, etc.) | N/A |
| | Complete thermocycling parameters | 95°C for 10 min followed by 50 cycles of 95°C for 20 s and 60°C for 40 sec |
| | Manufacturer of qPCR instrument | QuantStudio5 (Applied Biosystems) |
| | Specificity using genomic DNA (gel, sequence, melt or digest) | Gel |
| | Specificity using eDNA samples (gel, sequence, melt or digest) | Specificity was confirmed by direct amplicon sequencing (dual direction Sanger sequencing) of eDNA samples |

| | | |
|---|---|---|
| qPCR validation | For SYBR Green I, Cq of the NTC | N/A |
| | Standard curves with slope and y–intercept | Slope: -3.759; y–intercept: 44.672 |
| | PCR efficiency calculated from slope | 76-86% |
| | $R^2$ of standard curve | 0.95 |
| | Linear dynamic range | 1–1000000 DNA copies/reaction |
| | Cq variation at lower limit (LOQ) | Cq mean:39.061; Cq Standard deviation: 0.774; Cq coefficient of variation: 0.0198 |
| | Evidence for LOD & LOQ | LOD was set at the lowest standard were amplification was achieved and LOQ was set at the lowest standard with with 95% or greater detection |
| | If multiplex, efficiency and LOD of each assay | N/A |
| Data analysis | qPCR analysis program (source, version) | Applied Biosystems QuantStudio Design & analysis software 2.6.0 |
| | Cq method determination | Baseline Threshold: Cq is calculated using the PCR cycle number at which the fluorescence signal meets the threshold in the amplification plot. |
| | Outlier identification and disposition | Any wells that have Cq values that differ significantly (>Cq standard deviation with 95% confidence) from the average for the associated replicate wells. |
| | Results of NTCs | No amplification |
| | Justification of number and choice of reference genes | N/A |
| | Description of normalisation method | Standard curve |
| | Number and stage (RT or qPCR) of technical replicates | Six technical qPCR replicates of each biological replicate |
| | Repeatability (intra-assay variation) | $\Delta Ct \leq 5$ |

| Item to check | | Agra2 eDNA assay |
|---|---|---|
| Experimental design | Definition of experimental and control groups | Experimental: (1) genomic DNA (gDNA) from the target species<br>(2) environmental DNA (eDNA) samples consisting of soil and water from known infestation areas<br>Control: MilliQ water |
| | Number within each group | gDNA: n = 25; eDNA: n soil = 34; eDNA: n water = 20 |
| Sample (gDNA) | Description | Water samples were collected from four creeks/rivers adjacent to yellow crazy ant incursions. Soil samples were collected from three sites of known incursions. Tissue was sourced from five separate incursions in Cairns |
| | Microdissection or macrodissection | DNA from whole yellow crazy ant specimens was extracted |
| | Processing procedure:<br>If fixed – with what, how quickly? | Tissue samples were preserved immediately in 95% ethanol |
| | Sample storage condition and duration (especially for FFPE samples) | Extracted DNA was stored at 4°C |
| Sample (eDNA) | Description | Water samples: Direct collection and preservation of 30 mL water from water body adjacent to yellow crazy ant incursions<br>Soil samples: Direct collection of 1mL soil from sites of known incursions kept in ice for 2 hours and then extracted in the laboratory |
| | Processing procedure | Water samples were extracted using the PPLPP method; soil samples were extracted using the M1™ Bulk Sample Prep Kit for DNA – High Concentration (Biomeme, Inc), CTAB, PPLPP, Qiagen and Mu-DNA methods |
| | Sample storage condition and duration (especially for FFPE samples) | Extracted DNA was stored at 4°C for one week (until qPCR was performed) and subsequently stored at -20°C |
| Nucleic acid extraction (gDNA) | Procedure and/or instrumentation:<br>Name of kit and details of any modifications | Cetyltrimethylammonium bromide (CTAB) method |
| | Details of DNase or RNase | N/A |
| | Contamination assessment (DNA or RNA) | Nanodrop |

| | | |
|---|---|---|
| | Nucleic acid quantification | |
| | Instrument and method | Quantus ONE kit |
| | RNA integrity method/instrument | |
| | RIN/RQI or Cq of 3' and 5' transcripts | N/A |
| | Inhibition testing (Cq dilutions, spike or other) | N/A |
| Nucleic acid extraction (eDNA) | Procedure and/or instrumentation | |
| | Name of kit and details of any modifications | eDNA extarction methods: (1) PPLPP; (2) M1™ Bulk Sample Prep Kit for DNA – High Concentration, Biomeme, Inc; (3) CTAB; (4) modular-universal DNA extraction (Mu-DNA); (5) Qiagen® DNeasy PowerSoil kit. There was a subsequent purification using the DNeasy PowerClean CleanUp Kit (Qiagen®) |
| | Details of DNase or RNase | N/A |
| | Contamination assessment (DNA or RNA) | Nanodrop |
| | Nucleic acid quantification | |
| | Instrument and method | N/A |
| | RNA integrity method/instrument | |
| | RIN/RQI or Cq of 3' and 5' transcripts | N/A |
| | Inhibition testing (Cq dilutions, spike or other) | |
| qPCR target information | Gene symbol | N/A |
| | Sequence accession number | NCBI MZ330820-MZ330832 |
| | Amplicon length | 131 bp |
| | *In silico* specificity Screen (BLAST, etc.) | BLAST |

| | | |
|---|---|---|
| | Location of each primer and probe by exon or intron (if applicable) | |
| | What splice variants are targeted? | N/A |
| | Primer and probe sequences | Forward primer: 5'–ACTTTTGGTGCTTTAGGAATAATCTATGC–3'<br>Reverse primer: 5'–ATTGTTGCAGAAGTAAAATAAGCTCGA–3'<br>Probe: 5'–FAM-AGGCTTTATTGTTTGGGCTCATCACA-QSY–3' |
| | Location and identity of any modifications | Cytochrome Oxidase 1 (COI) region of the mitochondrial genome |
| | Complete reaction conditions: | |
| | Reaction volume and amount of cDNA/DNA | Reaction volume: 25 µL; amount of DNA: 5 µL |
| qPCR oligonucleotides | Primer, probe, $Mg^{2+}$ and dNTP concentrations | Each 25 µL reaction contained: 10 µL Environmental master Mix 2.0; 1 µL forward primer (10 µM); 1 µL reverse primer (10 µM); 1 µL probe (10 µM); 7 µL MilliQ water; 5 µL template DNA |
| | Polymerase identity and concentration | Applied Biosystems® TaqMan® Environmental Master Mix 2.0 (Thermo Fisher Scientific) |
| qPCR protocol | Buffer/kit identity and manufacturer | Applied Biosystems® TaqMan® Environmental Master Mix 2.0 (Thermo Fisher Scientific) |
| | Additives (SYBR Green I, DMSO, etc.) | N/A |
| | Complete thermocycling parameters | 95°C for 10 min followed by 50 cycles of 95°C for 20 s and 60°C for 40 sec |
| | Manufacturer of qPCR instrument | QuantStudio5 (Applied Biosystems) |
| | Specificity using genomic DNA (gel, sequence, melt or digest) | Gel |
| | Specificity using eDNA samples (gel, sequence, melt or digest) | Specificity was confirmed by direct amplicon sequencing (dual direction Sanger sequencing) of eDNA samples |
| | For SYBR Green I, Cq of the NTC | N/A |

| | | |
|---|---|---|
| qPCR validation | Standard curves with slope and y–intercept | Slope=-4.504, Y-intercept:50.362 |
| | PCR efficiency calculated from slope | 81-94% |
| | $R^2$ of standard curve | 0.997 |
| | Linear dynamic range | 1–1000000 DNA copies/reaction |
| | Cq variation at lower limit (LOQ) | Cq mean:37.214 ; Cq Standard deviation: 1.294; Cq coefficient of variation: 0.034 |
| | Evidence for LOD & LOQ | LOD was set at the lowest standard were amplification was achieved and LOQ was set at the lowest standard with with 95% or greater detection |
| | If multiplex, efficiency and LOD of each assay | N/A |
| Data analysis | qPCR analysis program (source, version) | Applied Biosystems QuantStudio Design & analysis software 2.6.0 |
| | Cq method determination | Baseline Threshold: Cq is calculated using the PCR cycle number at which the fluorescence signal meets the threshold in the amplification plot. |
| | Outlier identification and disposition | Any wells that have Cq values that differ significantly (>Cq standard deviation with 95% confidence) from the average for the associated replicate wells. |
| | Results of NTCs | No amplification |
| | Justification of number and choice of reference genes | N/A |
| | Description of normalisation method | Standard curve |
| | Number and stage (RT or qPCR) of technical replicates | Six technical qPCR replicates of each biological replicate |
| | Repeatability (intra-assay variation) | $\Delta Ct \leq 5$ |
